# Design of multi epitope-based peptide vaccine against E protein of human COVID-19: An immunoinformatics approach

**DOI:** 10.1101/2020.02.04.934232

**Authors:** Miyssa I. Abdelmageed, Abdelrahman H. Abdelmoneim, Mujahed I. Mustafa, Nafisa M. Elfadol, Naseem S. Murshed, Shaza W. Shantier, Abdelrafie M. Makhawi

**Author notes:** Corresponding author: Mujahed I. Mustafa.

## Abstract

**Background:** New endemic disease has been spread across Wuhan City, China on December 2019. Within few weeks, the World Health Organization (WHO) announced a novel coronavirus designated as coronavirus disease 2019 (COVID-19). In late January 2020, WHO declared the outbreak of a “public-health emergency of international concern” due to the rapid and increasing spread of the disease worldwide. Currently, there is no vaccine or approved treatment for this emerging infection; thus the objective of this study is to design a multi epitope peptide vaccine against COVID-19 using immunoinformatics approach.

**Method:** Several techniques facilitating the combination of immunoinformatics approach and comparative genomic approach were used in order to determine the potential peptides for designing the T cell epitopes-based peptide vaccine using the envelope protein of 2019-nCoV as a target.

**Results:** Extensive mutations, insertion and deletion were discovered with comparative sequencing in COVID-19 strain. Additionally, ten peptides binding to MHC class I and MHC class II were found to be promising candidates for vaccine design with adequate world population coverage of 88.5% and 99.99%, respectively.

**Conclusion:** T cell epitopes-based peptide vaccine was designed for COVID-19 using envelope protein as an immunogenic target. Nevertheless, the proposed vaccine is rapidly needed to be validated clinically in order to ensure its safety, immunogenic profile and to help on stopping this epidemic before it leads to devastating global outbreaks.

## 1. Introduction

Coronaviruses (CoV) are a large family of zoonotic viruses that cause illness ranging from the common cold to more severe diseases such as Middle East Respiratory Syndrome (MERS-CoV) and severe Acute Respiratory Syndrome (SARS-CoV). In the last decades, six strains of coronaviruses were identified, however in December 2019; a new strain has been spread across Wuhan City, China [1, 2]. It was designated as coronavirus disease 2019 (COVID-19) by the World Health Organization (WHO) [3]. In late January 2020, WHO declared the outbreak a global pandemic with cases in more than 45 countries where the COVID-19 spreading fast outside China, most significantly in South Korea, Italy and Iran with over 2,924 deaths and 85,212 cases confirmed while 39,537 recovered at 29 February 2020, 06:05 AM (GMT).

COVID-19 is a positive-sense single stranded RNA virus (+ssRNA). Its RNA sequence is approximately 30,000 bases in length [4]. It belongs to the subgenus *Sarbecovirus*, Genus *Betacoronavirus* within the family *Coronaviridae*. The corona envelope (E) protein is a small, integral membrane protein involved in several aspects of the virus’ life cycle, such as pathogenesis, envelope formation, assembly and budding; alongside with its interactions with both other CoVs proteins (M, N & S) and host cell proteins (release of infectious particles after budding) [5–9].

The infected person characterized with fever, upper or lower respiratory tract symptoms, or diarrhoea, lymphopenia, thrombocytopenia, and increased C-reactive protein and lactate dehydrogenase levels or combination of all these within 3-6 days after exposure. Further molecular diagnosis can be made by Real Time-PCR for genes encoding the internal RNA-dependent RNA polymerase and Spike’s receptor binding domain, which can be confirmed by Sanger sequencing and full genome analysis by NGS, multiplex nucleic acid amplification and microarray-based assays [10–14]

A phylogenetic tree of the mutation history of a family of viruses is possible to reconstruct with a sufficient number of sequenced genomes. The phylogenetic analysis indicates that COVID-19 is likely originated from bats [15]. It also showed that is highly related with at most seven mutations relative to a common ancestor [16].

The sequence of COVID-19 RBD, together with its RBM that contacts receptor angiotensin-converting enzyme 2 (ACE2), was found similar to that of SARS coronavirus. On January 2020, a group of scientists demonstrated that ACE2 could act as the receptor for COVID-19 [17–21].

However, COVID-19 differs from other previous strains in having several critical residues at 2019-nCoV receptor-binding motif (particularly Gln493) which provide advantageous interactions with human ACE2 [15]. This difference in affinity possibly explains why the novel coronavirus is more contagious than those other viruses.

At present, there is no vaccine or approved treatment for humans, but Chinese traditional medicine, such ShuFengJieDu Capsules and Lianhuaqingwen Capsule, could be possible treatments for COVID-19. However, there are no clinical trials approving the safety and efficacy for these drugs [22].

The main concept within all the immunizations is the ability of the vaccine to initiate an immune response in a faster mode than the pathogen itself. Although traditional vaccines, which depend on biochemical trials, induced potent neutralizing and protective responses in the immunized animals but they can be costly, allergenic, time consuming and require in vitro culture of pathogenic viruses leading to serious concern of safety [23, 24]. Thus the need for safe and efficacious vaccines is highly recommended.

Peptide-based vaccines do not need in vitro culture making them biologically safe, and their selectivity allows accurate activation of immune responses [25, 26]. The core mechanism of the peptide vaccines is built on the chemical method to synthesize the recognized B-cell and T-cell epitopes that are immunodominant and can induce specific immune responses. B-cell epitope of a target molecule can be linked with a T-cell epitope to make it immunogenic. The T-cell epitopes are short peptide fragments (8-20 amino acids), whereas the B-cell epitopes can be proteins [27, 28]. Therefore, in this study, we aimed to design a peptide-based vaccine to predict epitopes from corona envelope (E) protein using immunoinformatics analysis [29–34]. While rapid further studies are recommended to prove the efficiency of the predicted epitopes as a peptide vaccine against this emerging infection.

## 2. Materials and Methods

Workflow summarizing the procedures for the epitope-based peptide vaccine prediction is shown in **Fig. 1**

**Figure 1:**
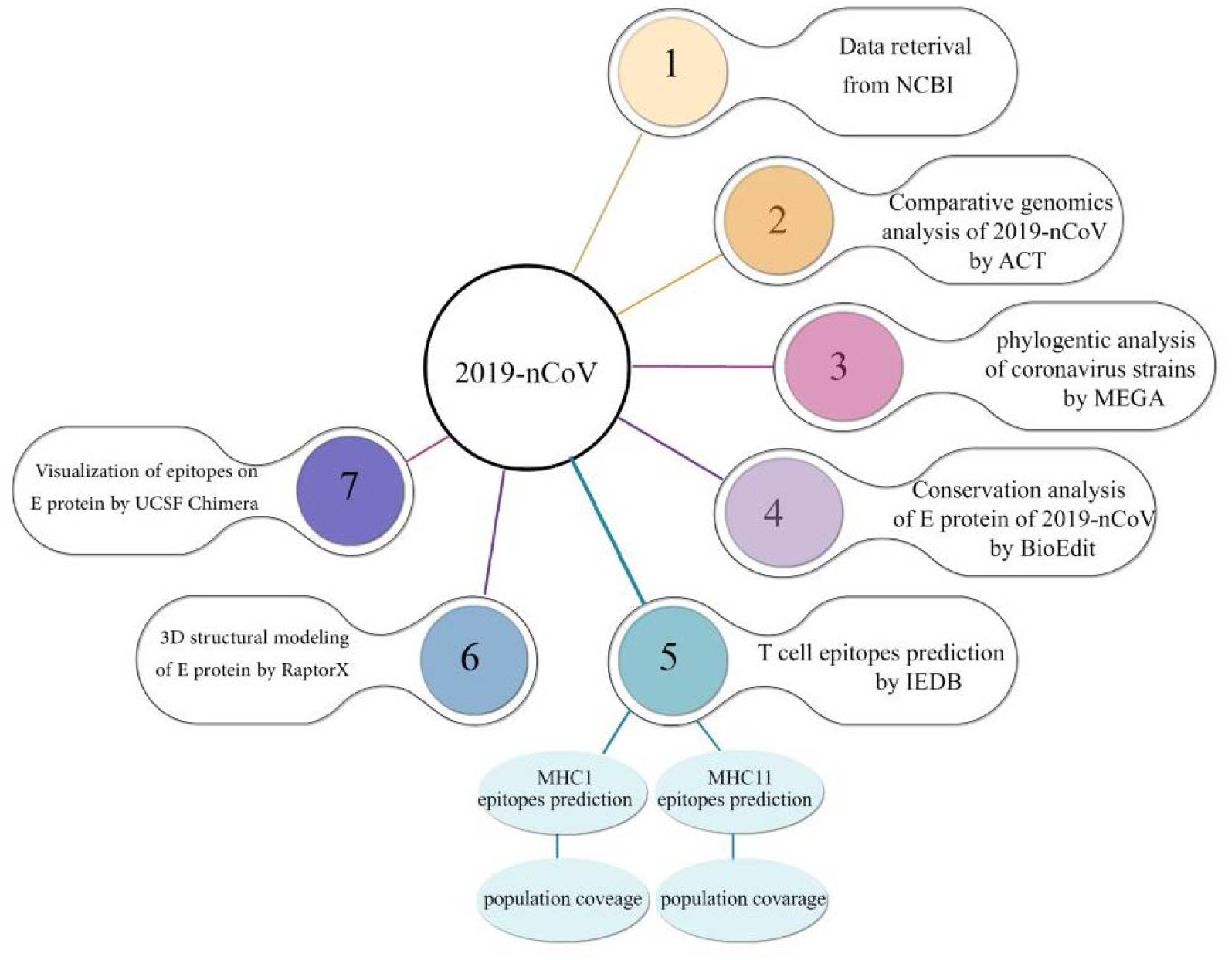
Descriptive workflow for the epitope-based peptide vaccine prediction.

### 2.1. Data retrieval

Full genebank files of the complete genomes and annotation of COVID-19(NC_04551) SARS-CoV(FJ211859), MESA-CoV(NC_019843), HCoV-HKU1(AY884001), (HCoV-OC43 (KF923903), HCoV-NL63 (NC_005831) and HCoV-229E (KY983587) were retrieved from the National Center of Biotechnology Information (NCBI); while The FASTA format of envelope (E) protein (YP_009724392.1), spike (S)protein (YP_009724390.1), nucleocapsid (N) protein (YP_009724397.2) and membrane (M) protein (YP_009724393.1) of 2019-nCoV and the envelope (E) protein of two Chinese and two American sequences (YP009724392.1, QHQ71975.1, QHO60596.1 and QHN73797.1) were obtained from the NCBI. (https://www.ncbi.nlm.nih.gov/)

### 2.2. The Artemis Comparison Tool (ACT)

ACT is an in silico analysis software for visualization of comparisons between complete genome sequences and associated annotations [35]. It is also applied to identify regions of similarity, rearrangements and insertions at any level from base-pair differences to the whole genome. (https://www.sanger.ac.uk/science/tools/artemis-comparison-tool-act).

### 2.3. VaxiJen server

It is the first server for alignment-independent prediction of protective antigens. It allows antigen classification solely based on the physicochemical properties of proteins without recourse to sequence alignment. It predicts the probability of the antigenicity one or multiple of protein based on auto cross covariance (ACC) transformation of protein sequence. Structural CoV-2019 protein (N,S,E and M) was analyzed by VaxiJen with threshold of 0.4 [36]. (http://www.ddg-pharmfac.net/vaxijen/VaxiJen/VaxiJen.html)

### 2.4. BioEdit

It is a software package proposed to stream a distinct program that can run nearly any sequence operation as well as a few basic alignment investigations. The sequences of E protein were retrieved from UniProt were run in BioEdit to determine the conserved sites through ClustalW in the application settings [37].

### 2.5. The Molecular Evolutionary Genetics Analysis (MEGA)

MEGA (version 10.1.6) is software for the comparative analysis of molecular sequences. It is used for pairwise and multiple sequences alignment alongside construction and analysis of phylogenetic trees and evolutionary relationships. The gap penalty was 15 for opening and 6.66 for extending the gap for both pairwise and multiple sequences alignment. Bootstrapping of 300 was used in construction of maximum like hood phylogenetic tree [38, 39]. (https://www.megasoftware.net).

### 2.6. Prediction of T-cell epitopes

IEDB tools were used to predict the conserved sequences (10-mersequence) from HLA class I and class II T-cell epitopes by using artificial neural network (ANN) approach [40–42]. Artificial Neural Network (ANN) version 2.2 was chosen as Prediction method as it depends on the median inhibitory concentration (IC50) [40, 43–45]. For the binding analysis, all the alleles were carefully chosen, and the length was set at 10 before prediction was done. Analysis of epitopes binding to MHC class I and II molecules was assessed by the IEDB MHC prediction server at (http://tools.iedb.org/mhci/) and (http://tools.iedb.org/mhcii/), respectively. All conserved immunodominant peptides binding to MHC I and II molecules at score equal or less than 100 median inhibitory concentrations (IC50) and 1000, respectively were selected for further analysis while epitopes with IC50 greater than 100 were eliminated [46].

### 2.7. Population coverage analysis

Population coverage for each epitope was carefully determined by the IEDB population coverage calculation tool. Due to the diverse binding sites of epitopes with different HLA allele, the most promising epitope candidates were calculated for population coverage against the whole world, China and Europe population to get and ensure a universal vaccine [47, 48]. (http://tools.iedb.org/population/)

### 2.8. Tertiary structure (3D) Modeling

The reference sequence of E protein that has been retrieved from gene bank was used as an input in RaptorX to predict the 3D structure of E protein [49, 50], the visualization of the obtained 3D protein structure was performed in UCSF Chimera (version1.8) [51].

### 2.9. In silico Molecular Docking

#### 2.9.1. Ligand Preparation

In order to estimate the binding affinities between the epitopes and molecular structure of MHC I and MHC II, in silico molecular docking were used. Sequences of proposed epitopes were selected from COVID-19 reference sequence using UCSF Chimera 1.10 and saved as a PDB file. The obtained files were then optimized and energy minimized. The HLA-A*02:01 was selected as the macromolecule for docking. Its crystal structure (4UQ3) was downloaded from the RCSB Protein Data Bank (http://www.rcsb.org/pdb/home/home.do), which was in a complex with an azobenzene-containing peptide [52].

All water molecules and heteroatoms in the retrieved target file 4UQ3 were then removed. Target structure was further optimized and energy minimized using Swiss PDB viewer V.4.1.0 software[53].

#### 2.9.2. Molecular docking

Molecular docking was performed using Autodock 4.0 software, based on Lamarckian Genetic Algorithm; which combines energy evaluation through grids of affinity potential to find the suitable binding position for a ligand on a given protein[54,55]Polar hydrogen atoms were added to the protein targets and Kollman united atomic charges were computed. The target’s grid map was calculated and set to 60×60×60 points with grid spacing of 0.375 Å. The grid box was then allocated properly in the target to include the active residue in the center. The genetic algorithm and its run were set to 100.The docking algorithms were set to default. Finally, results were retrieved as binding energies and poses that showed lowest binding energies were visualized using UCSF Chimera.

## 3. Results

### 3.1. The Artemis Comparison Tool

The reference sequence of envelope protein was aligned with HCov-HKU1 reference protein using artemis comparison tool as illustrated in (Fig. 2).

**Figure 2:**
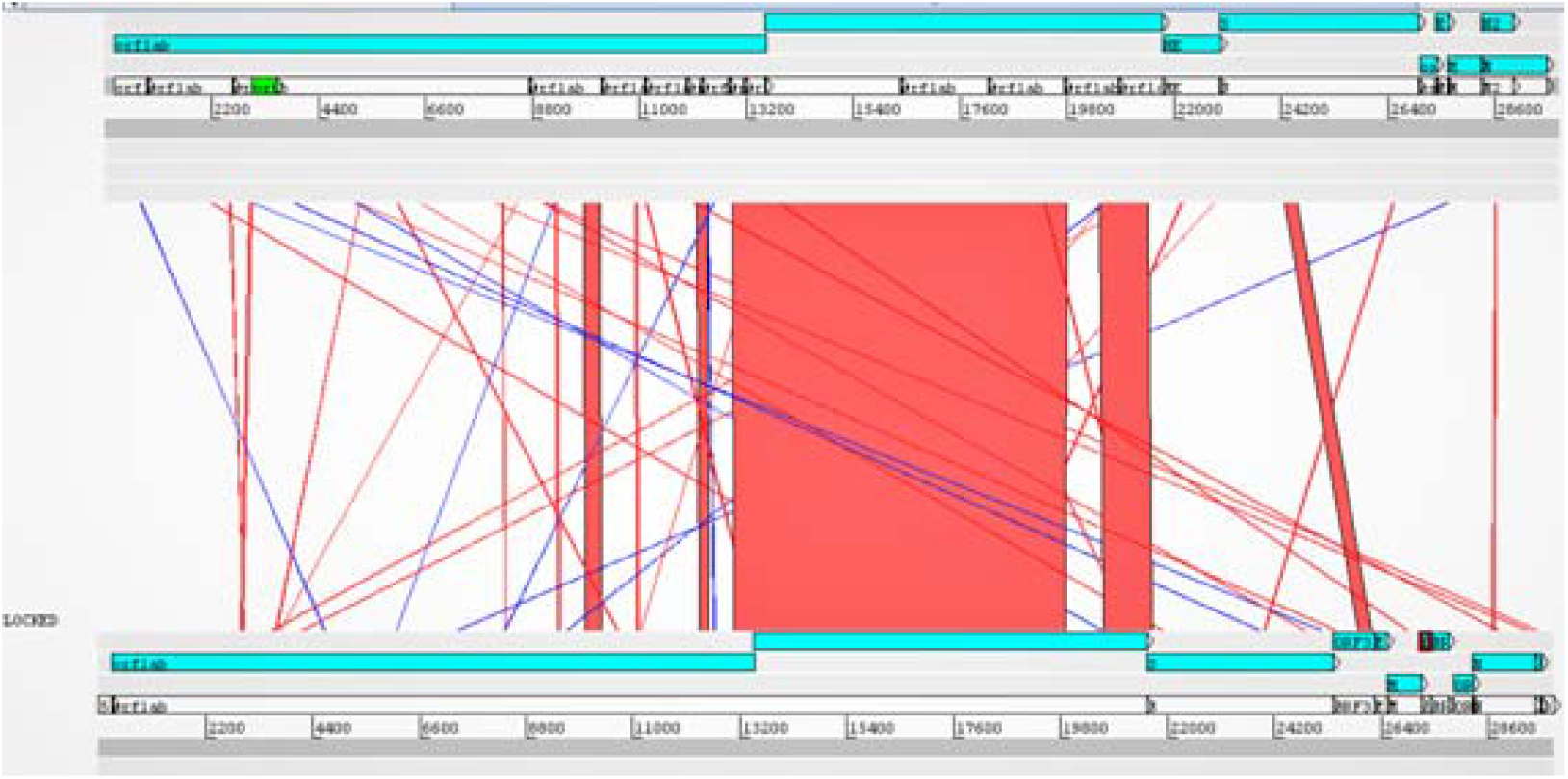
Artemis analysis of envelope protein displaying 3 windows, the upper window represents HCov-HKU1 reference sequence and its genes are highlighted in blue starting from *orflab* gene and ending with *N* gene. The middle window describes the similarities and the difference between the two genomes. Red lines indicate match between genes from the two genomes blue lines indicates inversion which represents same sequences in the two genomes but they are organized in the opposite direction, and the lower windows represents COVID-19and its genes started from *ARFLAB* and ends with *N* genes.

### 3.2. VaxiJen server

The mutated proteins were tested for antigenicity using VaxiJen software, where the envelope protein found as the best immunogenic target in Table 1.

**Table 1:**
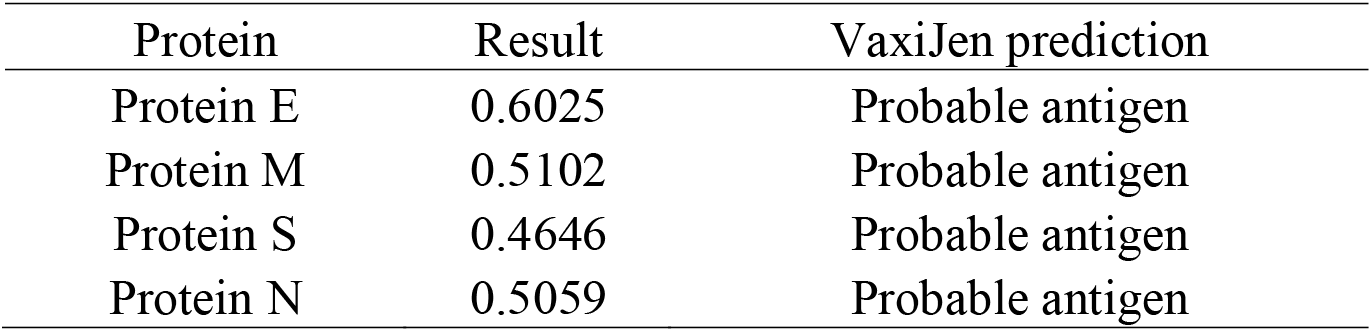
VaxiJen overall prediction of probable COVID-19 antigen:

| Protein | Result | VaxiJen prediction |
| --- | --- | --- |
| Protein E | 0.6025 | Probable antigen |
| Protein M | 0.5102 | Probable antigen |
| Protein S | 0.4646 | Probable antigen |
| Protein N | 0.5059 | Probable antigen |

### 3.3. BioEdit

Sequence alignment of COVID-19envelope protein was done using BioEdit software which shows total conservation across four sequences which were retrieved from China and USA (Fig. 3).

**Figure 3:**
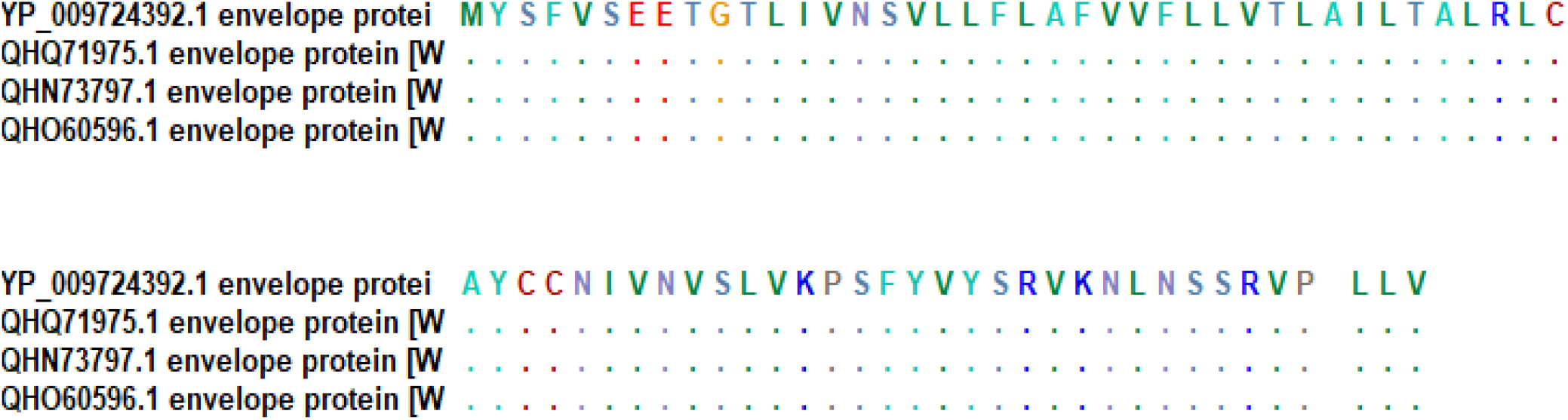
Sequence alignment of envelope protein of COVID-19 using BioEdit software (total conservation through the 4 strains: 2 form China and 2 from USA)

### 3.4. The Molecular Evolutionary Genetics Analysis

To study the evolutionary relationship between all the seven strains of coronavirus, a multiple sequence alignment (MSA) was performed using ClustalW by MEGA software. This alignment was used to construct maximum likelihood phylogenetic tree as seen in **Fig. 4**.

**Figure 4:**
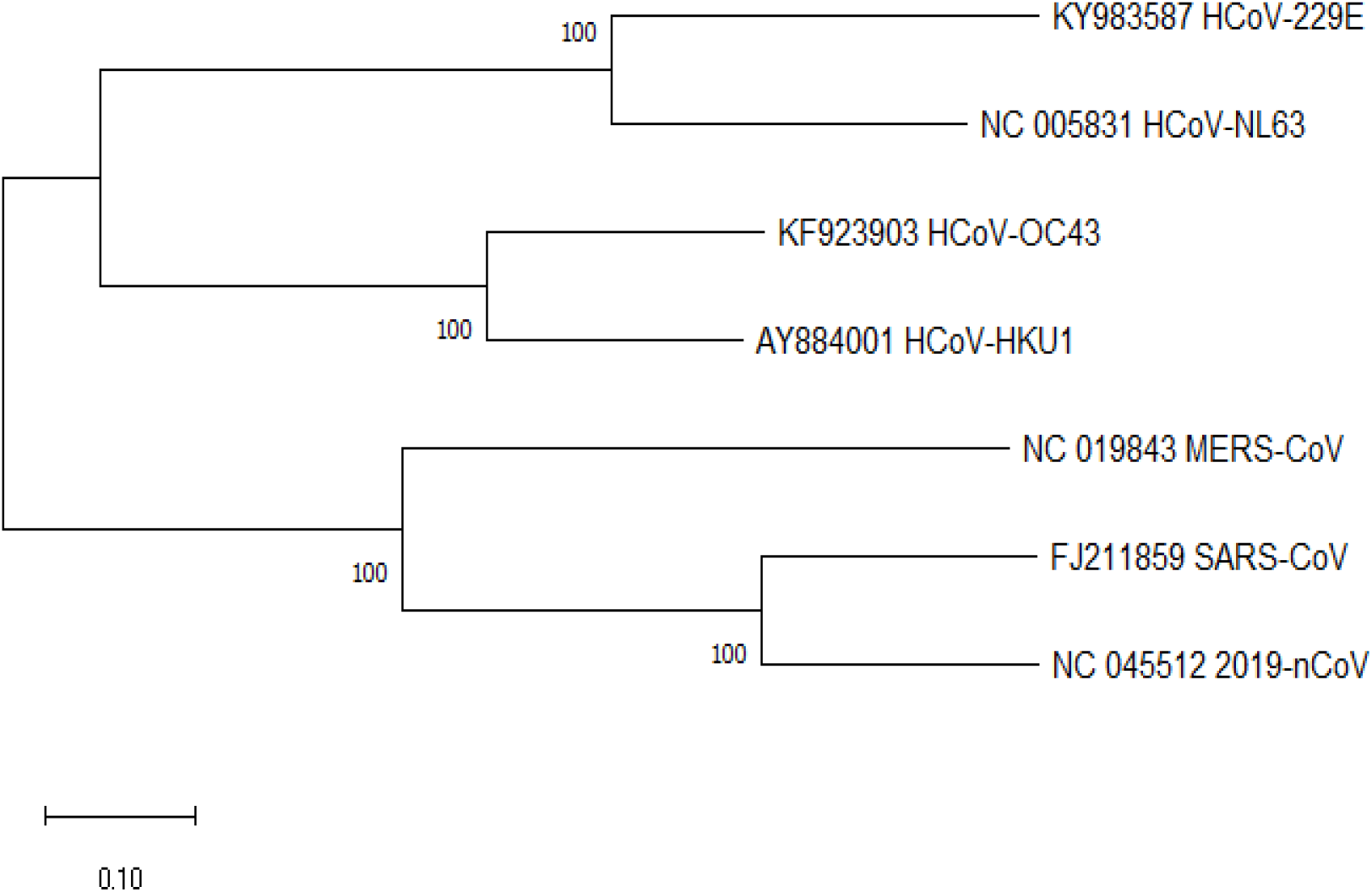
Maximum likelihood phylogenetic tree which describes the evolutionary relationship between the seven strains of coronavirus.

### 3.5. Prediction of T-cell epitopes and population coverage

IEDB website was used to analyze 2019-nCoV envelope protein for T cell related peptides. Results show ten MHC class I and II associated peptides with high population coverage (Tables 2 and 3; Fig. 5). The most promising peptides were visualized using UCSF Chimera software (Fig.6a and b).

**Figure 5:**
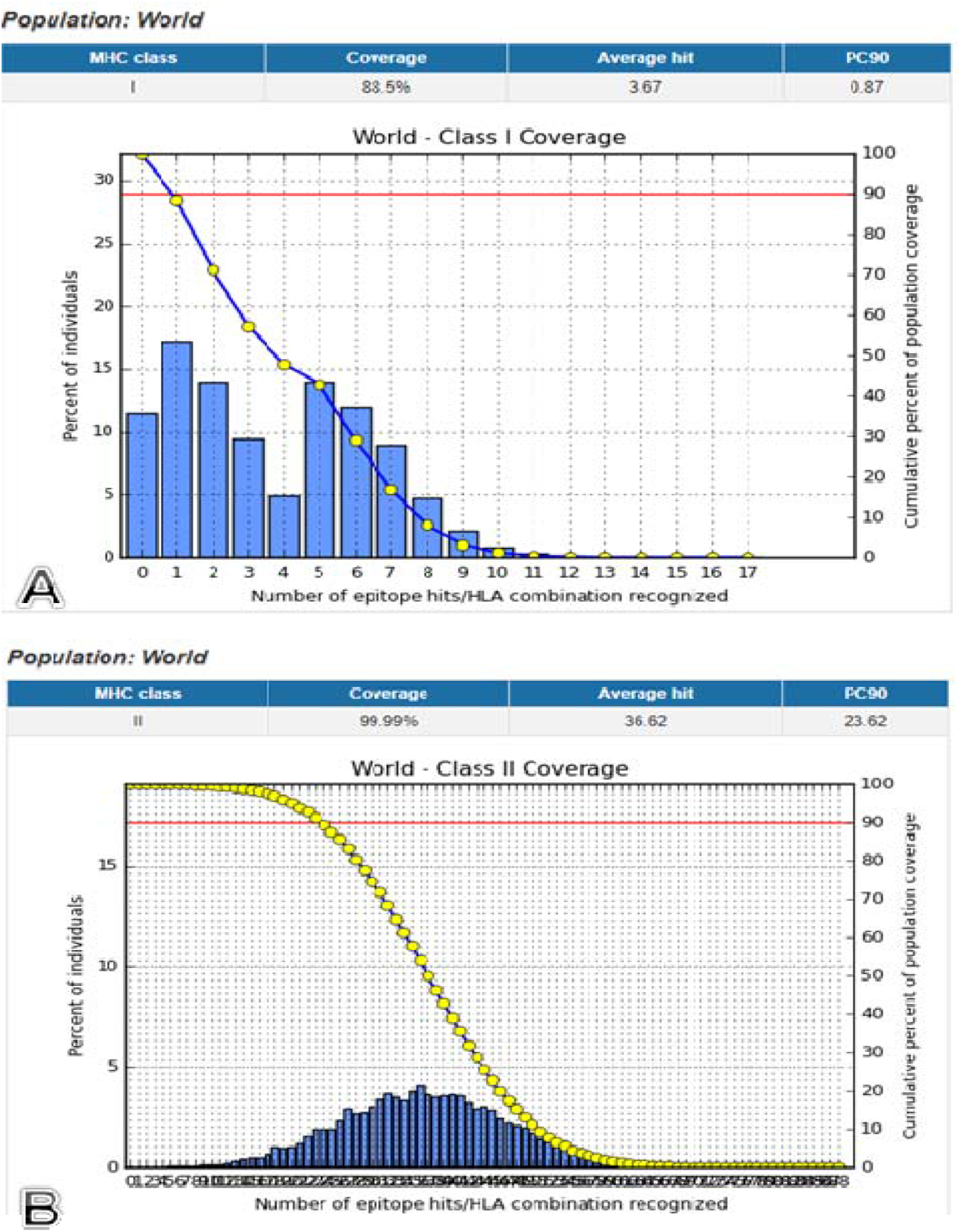
Schematic diagrams (**A**) and (**B**) showing world population coverage of envelope protein of COVID-19 binding to MHC class I and MHC class II molecule respectively.

**Figure 6:**
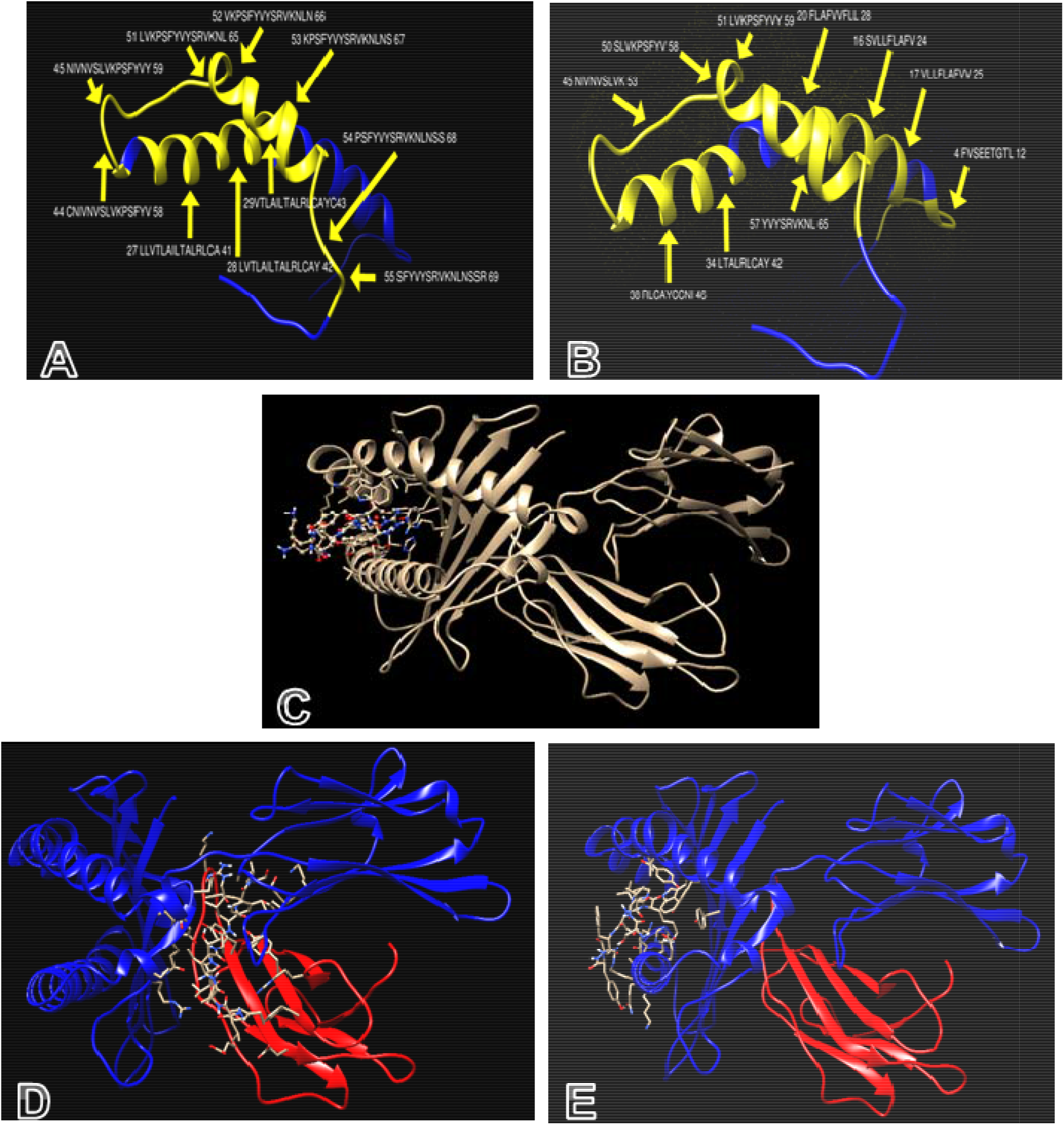
3D structures visualized by UCSF Chimera: (**A**) and (**B**) showing the most promising peptides in envelope protein of COVID-19 (yellow colored) binding to MHC class I and MHC class II respectively, while (**C, D**, and **E**) showing the molecular docking of YVYSRVKNL, LAILTALRL and SLVKPSFYV peptides of coronavirus docked in HLA-A*02:01 respectively

**Table 2:**
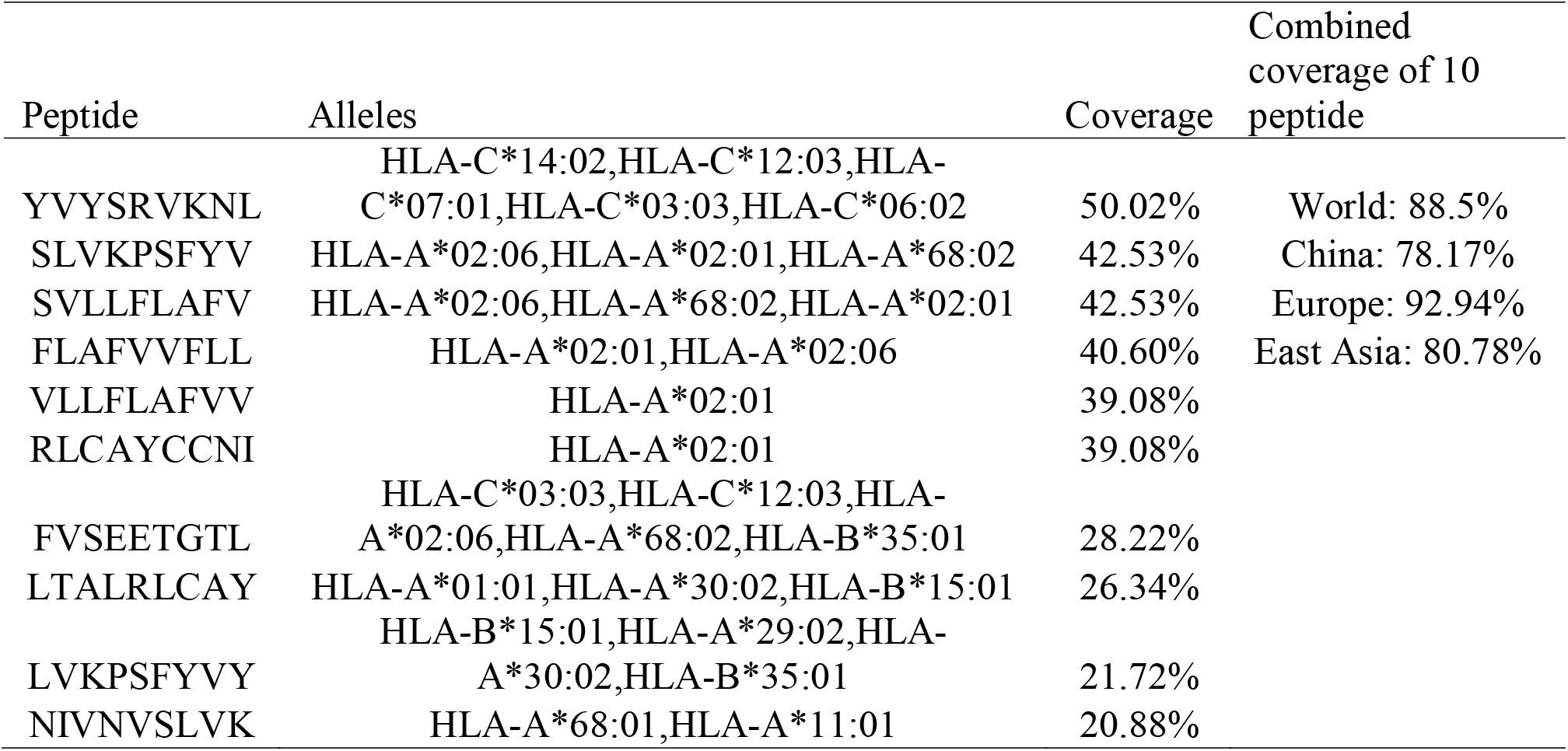
The most promised MHC class I related peptides in envelope protein based vaccine of COVID-19 along with the predicted World, China, Europe and East Asia coverage:

**Table 3:**
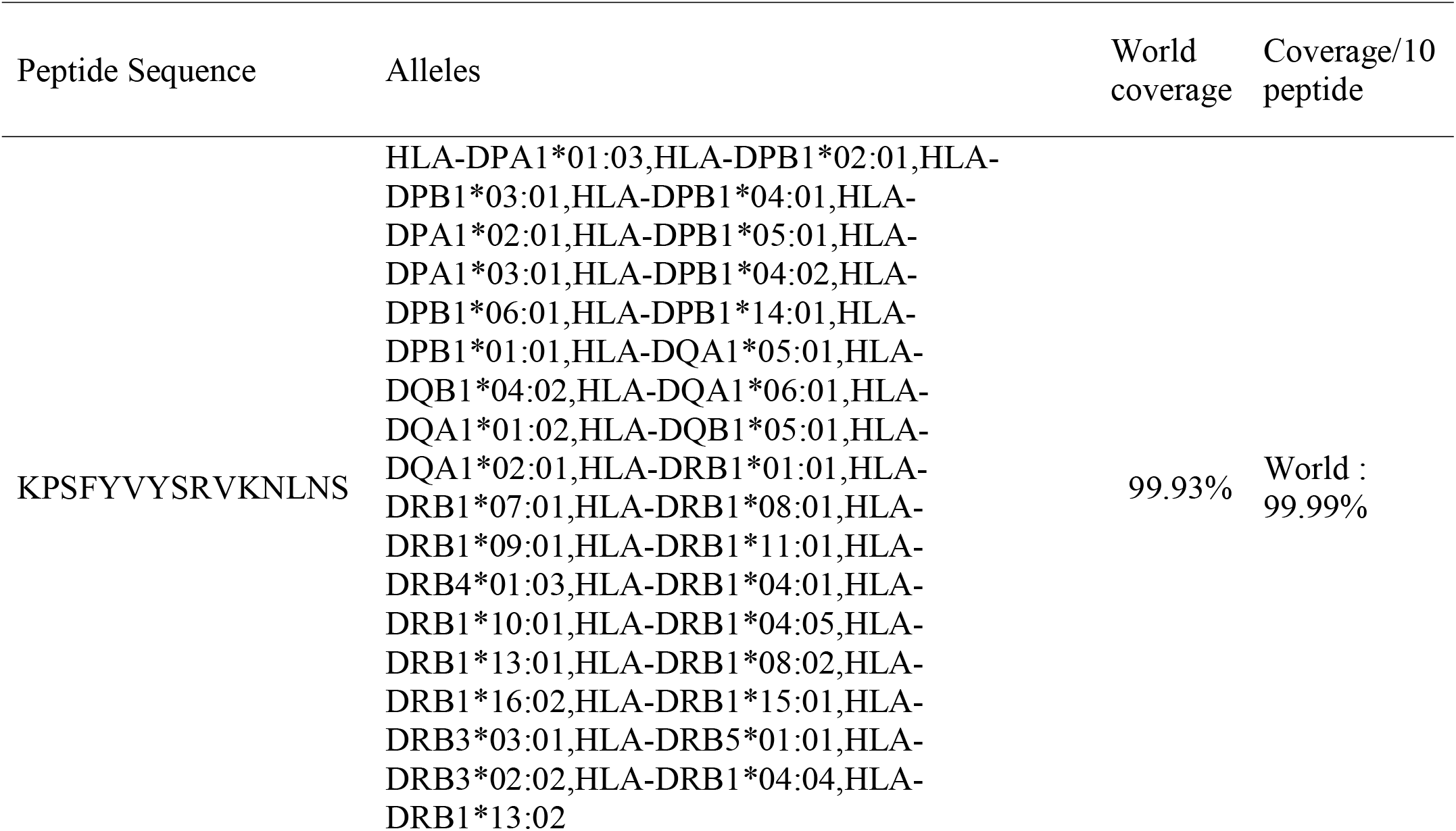
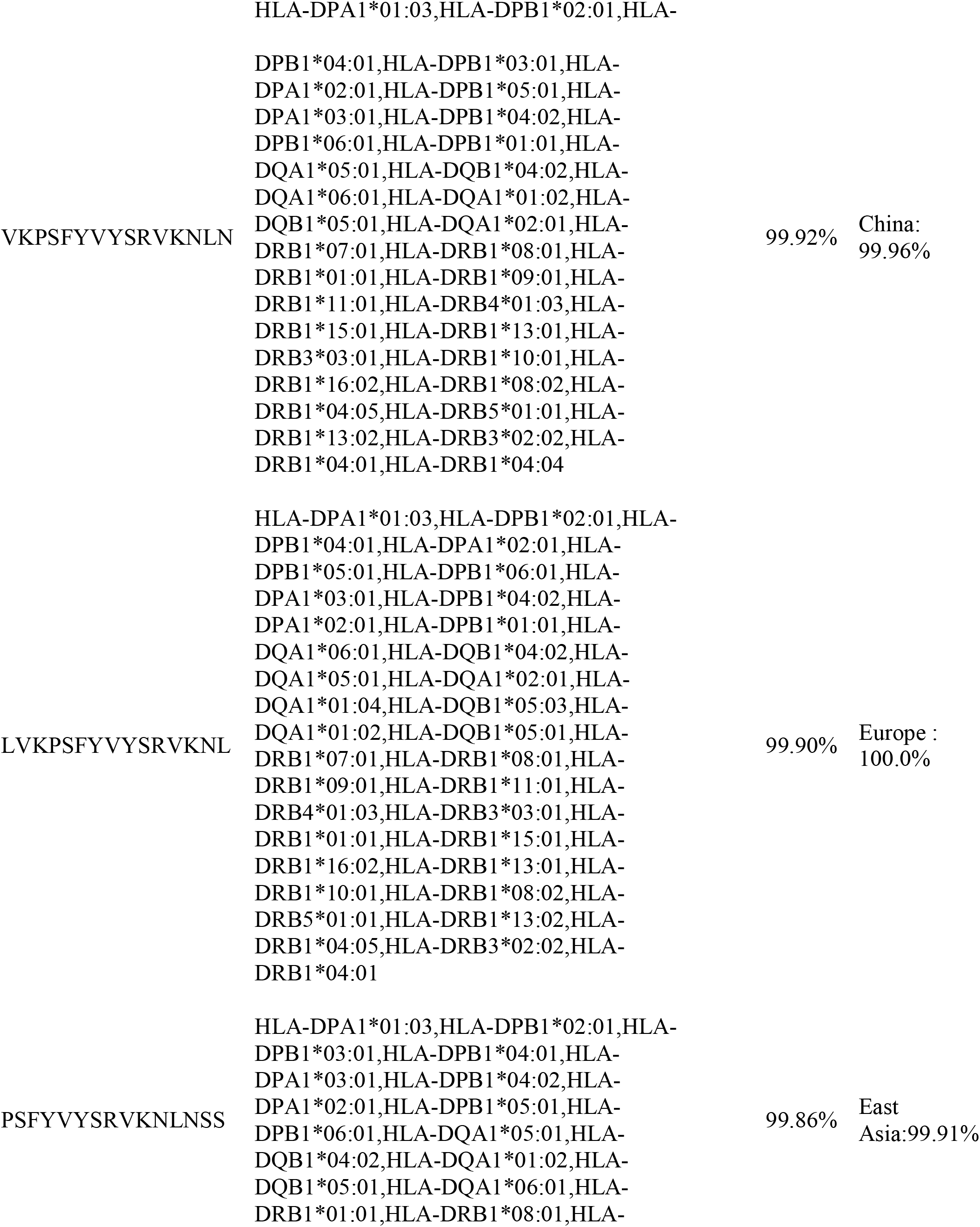
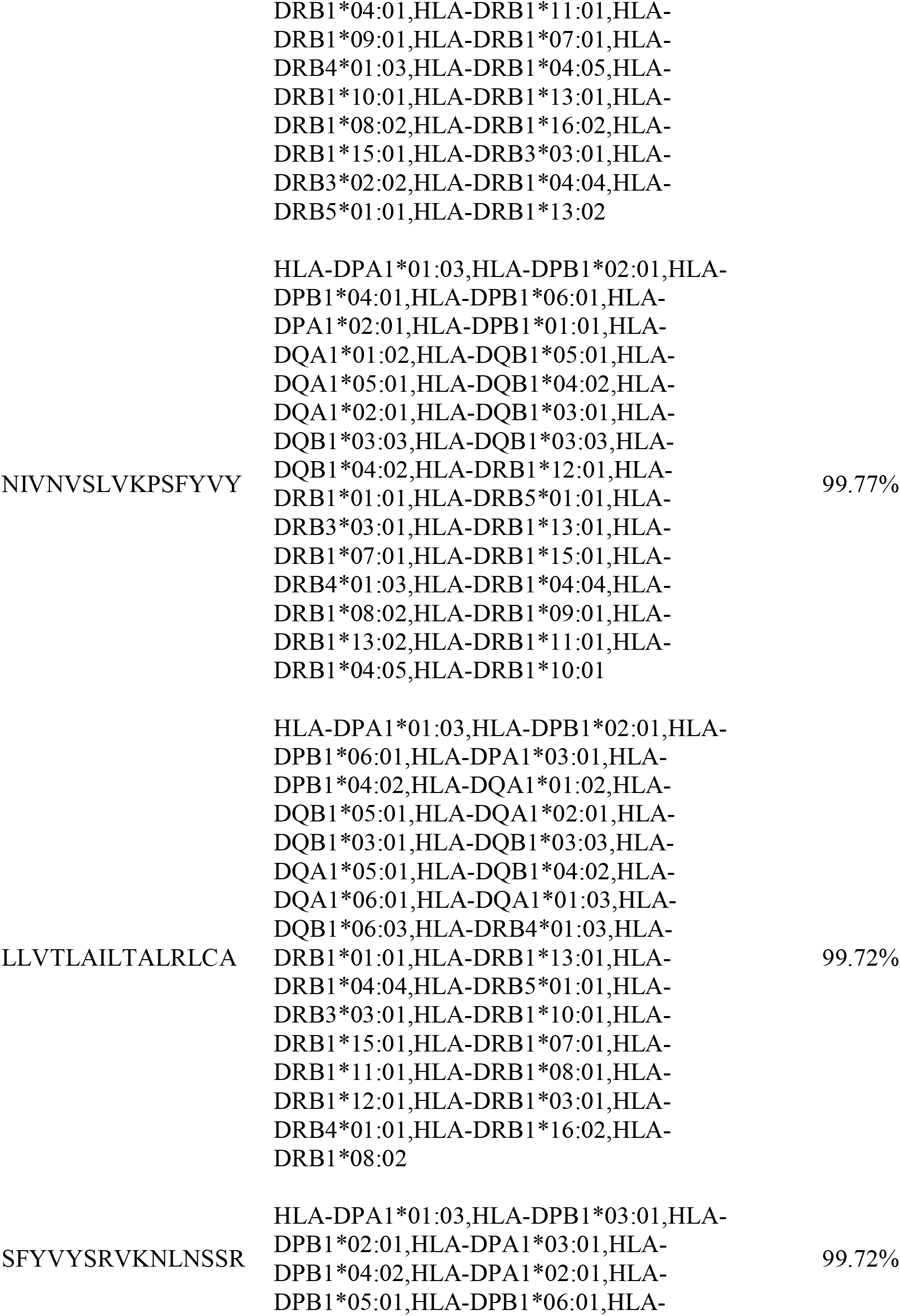
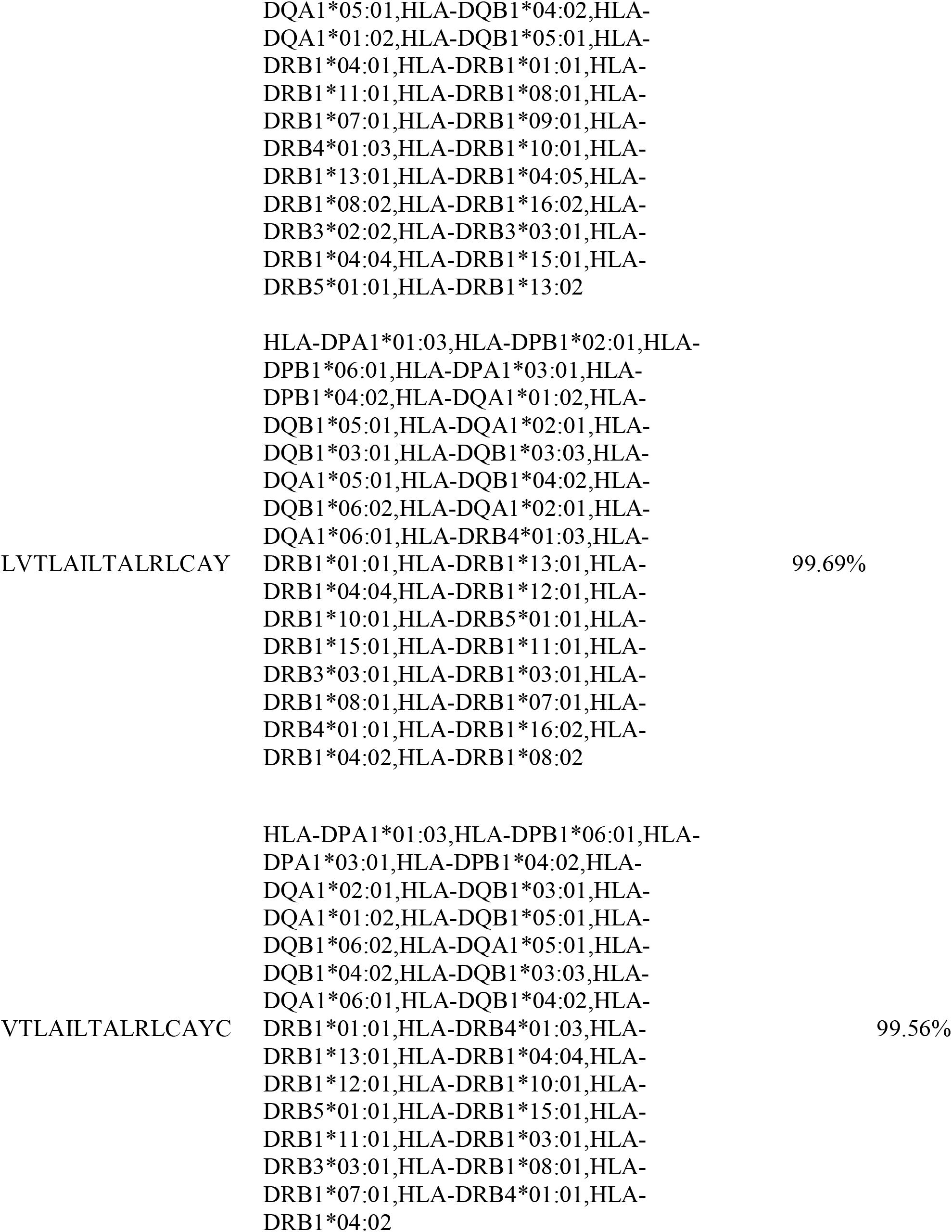
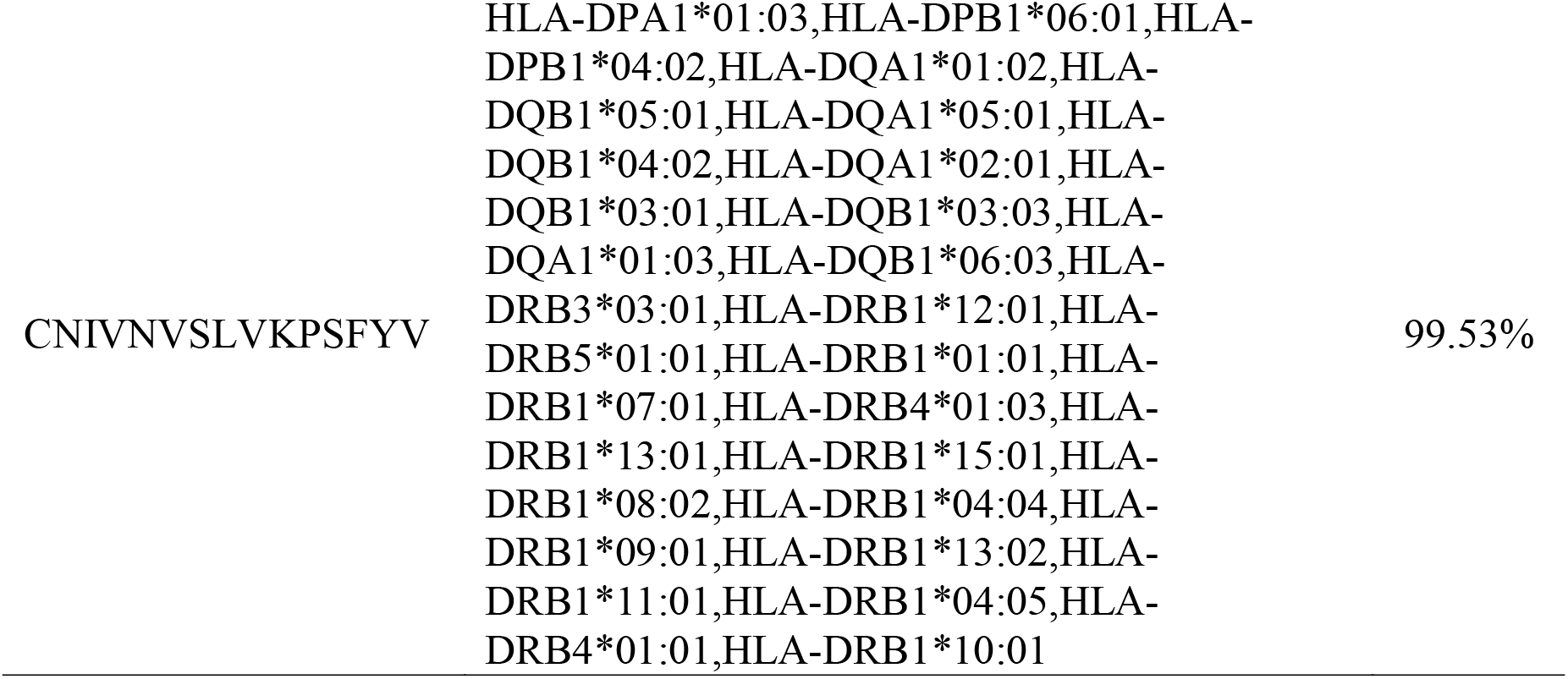
The most promised MHC class II related peptides in envelope protein based vaccine of COVID-19 along with the predicted World, China, Europe and East Asia coverage:

## 4. Discussion

Designing of a novel vaccine is very crucial to defending the rapid endless of global burden of disease [56–59]. In the last few decades, biotechnology has advanced rapidly; alongside with the understanding of immunology which assisted the rise of new approaches towards rational vaccines design [60]. Peptide-based vaccines are designed to elicit immunity particular pathogens by selectively stimulating antigen specific for B and T cells [61].Applying the advanced bioinformatics tools and databases, various peptide-based vaccines could be designed where the peptides act as ligands [62–64]. This approach has been used frequently in Saint Louis encephalitis virus [65], dengue virus [66], chikungunya virus [67] proposing promising peptides for designing vaccines.

The COVID-19 is an RNA virus which tends to mutate more commonly than the DNA viruses [68]. These mutations lied on the surface of the protein, which make COVID-19 more superior than other previous strains by inducing its sustainability leaving the immune system in blind spot [69].

In our present work, different peptides were proposed for the designing of vaccine against COVID-19 (Fig. 1). In the beginning, the whole genome of COVID-19 was analyzed by comparative genomic approach to determine the potential antigenic target [70]. Artemis Comparative Tool (ACT) was used to analyze human Coronavirus (HCov-HKU1) reference sequence vs. Wuhan-Hu-1 COVID-19. Results obtained (Fig.2) revealed extensive mutation among the tested genomes. New genes; *ORF8* and *ORF6*; were found inserted in COVID-19 which were absent in HCov-HKU1 that might be acquired by the horizontal gene transmission [71]. High rate of mutation between the two genomes were observed in the region from 20000 bp to the end of the sequence. This region encodes the four major structural proteins in coronavirus which are envelope (E) protein, nucleocapsid (N) protein, membrane (M) protein, and spike (S) protein, all of which are required to produce a structurally complete virus [72, 73].

These conserved antigenic sites were revealed in previous studies through sequence alignment between MERS-CoV and Bat-coronavirus [74] and analyzed in SARS-CoV [75].

The four proteins were then analyzed by Vaxigen software to test the probability of antigenic proteins. Protein E was found to be the most antigenic gene with the highest probability as shown in Table 1. Literature survey confirmed this result in which protein E was investigated in severe acute respiratory syndrome (SARS) in 2003 and, more recently, Middle-East respiratory syndrome (MERS) [72]. Furthermore, the conservation of this protein against the seven strains was tested and confirmed through the use of BioEdit package tool (Fig. 3).

As phylogenetic analysis is very powerful tool for determining the evolutionary relationship between strains. Multiple sequence alignment (MSA) was performed using ClustalW for the seven strains of coronavirus, which are COVID-19(NC_04551), SARS-CoV (FJ211859), MESA-CoV (NC_019843), HCoV-HKU1 (AY884001), HCoV-OC43 (KF923903), HCoV-NL63 (NC_005831) and HCoV-229E (KY983587). The maximum likelihood phylogenetic tree revealed that COVID-19 is found in the same clade of SARS-CoV, thus the two strains are highly related to each other (Fig. 4).

The immune response of T cell is considered as a long lasting response compared to B cell, where the antigen can easily escape the antibody memory response [76]. Vaccines that effectively generate cell-mediated response are needed to provide protection against the invading pathogen. Moreover the CD8+ T and CD4+ T cell responses play a major role in antiviral immunity [77]. Thus designing vaccine against T cell is much more important.

Choosing protein E as the antigenic site, the binding affinity to MHC molecules was then evaluated. The protein reference sequence was submitted to IEDB MHC predication tool. 21 peptides were found to bind MHC class I with different affinities (Table 1), from which ten peptides were selected for vaccine design based on the number of alleles and world population percentage (Table 2; Fig. 5).Analysis in IEDB MHC II binding prediction tool resulted in prediction of 61 peptides (Table 2), from which ten peptides were selected for vaccine design based on the number of alleles and world population percentage (Table 3; Fig.5). Unfortunately, IEDB did not give any result for B cell epitopes, this might be due to the length of the COVID-19 (75 amino acids).

It is well known that peptides recognized with high number of HLA molecules are potentially inducing immune response. Based on the aforementioned results and taking into consideration the high binding affinity to both MHC classes I and II, conservancy and population coverage, three peptides are strongly proposed to formulate a new vaccine against COVID-19.

These findings were further confirmed by the results obtained for the molecular docking of the proposed peptides and HLA-A*02:01. The formed complex between MHC molecule and the three peptides (YVYSRVKNL, SLVKPSFYV and LAILTALRL) have shown peptide amino-and carboxyl-termini forming one and three hydrogen bonds, respectively at the two ends of a binding groove with MHC residues with least binding energy −13.2 kcal/mol, −11 kcal/mol and −11.3 kcal/mol, respectively (Fig.6C, D and E).

Although both flu and anti-HIV drugs are used currently in China for treatment of COVID-19, and chloroquine phosphate, an old drug for treatment of malaria, has recently found to have apparent efficacy and acceptable safety against COVID-19 [78,79]; nevertheless more studies are required to standardize these therapies. In addition, there has been some success in the development of mouse models of MERS-CoV and SARS-CoV infection, and candidate vaccines where the envelope (E) protein is mutated or deleted have been described [80–86]. To best of our knowledge, this is the first study to identify certain peptides in envelope (E) protein as candidates for COVID-19. Accordingly, these epitopes were strongly recommended as promising epitopes vaccine candidate against T cell.

## 5. Conclusion

Extensive mutations, insertion and deletion were discovered in COVID-19 strain using the comparative sequencing. In addition, a number of MHC class I and II related peptides were found promising candidates. Among which the peptides YVYSRVKNL, SLVKPSFYV and LAILTALRL show high potentiality for vaccine design with adequate world population coverage. T cell epitope-based peptide vaccine was designed for COVID-19 using envelope protein as an immunogenic target; nevertheless, the proposed vaccine rapidly needs to be validated clinically ensuring its safety and immunogenic profile to help on stopping this epidemic before it leads to devastating global outbreaks.

## Acknowledgement

The authors acknowledge the Deanship of Scientific Research at University of Bahri for the supportive cooperation.

## Data Availability

All data underlying the results are available as part of the article, and no additional source data are required.

## Authors’ contributions

MIA: Conceptualization, Formal analysis, Investigation, Methodology, Validation, Visualization, Writing– original draft. AHA: Formal analysis, Investigation, Methodology. MIM: Methodology, Writing– original draft, Writing – review and editing. NME: Formal analysis, Methodology, Visualization. NSM: Conceptualization, Resources, Writing – review & editing. SWS: Visualization, Validation, Writing – review and editing, AMM: Data curation, Conceptualization, Project administration, Supervision, Writing - review and editing. All authors have read and approved the final manuscript.

## Abbreviations

WHO: World Health Organization
CoV: Coronaviruses
MERS-CoV: Middle East Respiratory Syndrome
SARS-CoV: Acute Respiratory Syndrome
COVID-19: novel coronavirus
HCoV-HKU1: Human coronavirus HKU1
HCoV-OC43: Human coronavirus OC43
HCoV-NL63: Human coronavirus NL63
HCoV-229E: Human coronavirus 229E
ACE2: angiotensin converting enzyme 2
RBM: receptor-binding motif
RBM: receptor-binding domain
vs: versus
ACT: Artemis Comparison Tool
ACC: auto cross covariance
MEGA: Molecular Evolutionary Genetics Analysis
ANN: Artificial Neural Network
IEDB: Immune Epitope Database
IC50: median inhibitory concentrations
MHC I: Major Histocompatibility complex class I
MHC II: Major Histocompatibility complex class II
PDB: Protein database
MSA: multiple sequence alignment

